# Different liposomal formulations of 5-Fluorouracil result in variations to gastrointestinal toxicities

**DOI:** 10.1101/2024.12.16.628814

**Authors:** Daniel Thorpe, Wen Wang, Robert Milne, Andrea Stringer

## Abstract

5-Fluorouracil (5FU) treatment can induce severe mucositis, gastrointestinal toxicity. New formulations of 5-FU increase cancer cytotoxicity, such as neutral liposomes (NL), cationic liposomes (CL), and polyethylenimine copper (PEI-Cu) complexes. However, gastrointestinal toxicity for these new formulations remains unclear. We aim to determine if new formulations in 5FU reduce mucositis by reducing intestinal goblet cell and nerve integrity.

Sprague Dawley rats were randomly assigned to groups: saline (n=6), 5FU (n=6 ), NL-5FU (n=5), CL-5FU (n=5), PEI-CU-5FU (n=6), NL-PEI-Cu-5FU (n=5) CL-PEI-Cu-5FU (n=5) (Formulations were equivalent to 10mg/kg 5FU in saline). Treatment was administered daily for 5 days. Rats where humanely killed 2 days after treatment. Haematoxylin and Eosin staining for histological change, Alcian Blue-PAS staining for mucin composition, and immunohistochemistry with S100 antibody for nerve integrity were performed. Statistical analysis using Kruskal-Wallis test with Dunns post-test and Mann Whitney U test d were performed. Effect size was determined using Cohen’s D test.

In the jejunum, inflammatory infiltrate increased in PEI-Cu-5FU rats compared to 5-FU controls (p=0.0011). S100 positive nerve bundles increased in NL-5FU rats compared to saline control (p<0.05). S100 positive cells increased in CL-PEI-Cu-5FU rats compared to saline controls.

PEI-Cu-5FU formulation was associated with increased inflammatory infiltrate potentially in response to damage in the jejunum. However, liposomal formulations increased S100 positive neural cells, which may offer protection through increased gastrointestinal motility and contraction. While these formulations of 5FU increase cancer cytotoxicity, the gastrointestinal toxicity remains similar. However, further close monitoring of 5FU formulations for gastrointestinal toxicity is warranted.

## Introduction

5-Fluorouracil (5FU) is a cytotoxic chemotherapeutic agent renowned for causing mucositis, with the most severe damage occurring in the small intestine. Carriers of 5FU such as liposomes, with 5FU entrapped as a complex with polyethylenimine copper (PEI-Cu), offer an increased plasma circulation time and the potential for an increase in the cytotoxicity for cancer cells [1-6]. While 5FU formulated into liposome has been found to increase cytotoxicity [1, 5], no studies have been undertaken to determine whether any toxic side effects may occur in the gut from administering liposomal formulations of 5FU as a complex with PEI-Cu.

Carriers of 5FU have increased efficacy in vitro models [4] and ex vivo models of cancer [2], and liposomal carriers of chemotherapy agents in, in vivo studies have shown increased tumour cytotoxicity [3, 7, 8]. Dupertuis et.al. demonstrated increased cytotoxicity against two human colorectal cancer cell lines LS174T and HT-29 when liposomes formulated with docosahexaenoic acid and eicosapentaenoic acid at a ratio of 1:2 were used to carry 5-FU. The liposomal formulations significantly increased apoptosis in HT-29 cells (P = 0.001), necrosis in LS174T (P = 0.02) and HT-29 cells (P = 0.004) respectively. Ahman et.al. also demonstrated a significant increase in cytotoxicity in SK-MEL-5 cancer cells when 5-FU-nanoemulsion (P < 0.05), and 5-FU-nanoemulsion-Gel (P < 0.01) was administered compared to 5-FU alone. This demonstrates that carriers of 5FU in in vitro models have increased cytotoxicity compared to 5-FU alone. Using an ex vivo model of excised cow, goat and rat skin, increased permeability and increased drug release were shown in 5-FU-nanoemulsion and 5-FU-nanoemulsion gel compared to 5-FU alone [2].. This increase in cytotoxicity is not limited to 5-FU chemotherapy agents Oxaliplatin Irinotecan and Paclitaxel when administered with a liposomal carrier also had increased cytotoxicity in vivo. Oxaliplatin (5 mg/kg i.v. on days 4, 8 and 12 post tumour inoculation) containing (polyethylene glycol) PEG-coated cationic liposomes significantly (P < 0.01) decreased tumour volume in C57BL/6 mice bearing the Lewis lung carcinoma cell model [8]. Irinotecan (5 mg/kg i.v. every 3 days) containing PEG-coated liposomes significantly prevented tumour growth in BALB/c mice with 4T1 xenograft tumours compared to Irinotecan alone [7]. Paclitaxel (1 mg/kg i.v. every 3 days) containing PEG-coated liposomes significantly (P < 0.0001) reduced tumour growth compared to Paclitaxel alone [9]. This suggests that chemotherapy delivery can have increased tumour cytotoxicity when administered with a carrier. 5FU-induced mucositis results in pathological change in the gastrointestinal tract associated with pain, ulceration, bloating, vomiting and diarrhoea. Histopathological change resulted in decreased epithelial surface area and therefore the area for absorption from the lumen into the body decreased, potentially increasing the likelihood of malnutrition [10]. Changes to mucin composition, increasing cavitated goblet cells and therefore depleting mucin stores which may result in a decrease to the protective capacity of the mucus barrier [10]. Finally, enteric nervous system loss has been observed [11] which may result in the alternations in mucin composition and mucus barrier loss. Therefore, this study aimed to determine if the use of liposomes and PEI-Cu with 5FU was associated with modifications to histopathology, mucin composition and neural structure during mucositis compared with standard 5FU.

## Methods

### Ethics

Approval for the use of animals was granted by SA pathology (41A.12). Rats were monitored two times daily, and if any rat showed a dull ruffled coat with accompanying dull and sunken eyes, was cold to touch with no spontaneous movement and a hunched appearance, then they were euthanized.

### Liposomal preparation

The polyethylenimine (Sigma-Aldrich, Australia) copper complex was prepared by adding polyethlylenimine to copper acetate (Sigma-Aldrich, Australia) at a ratio of 1:4 and stirred till a purple blue homogenous solution was obtained, for a final copper concentration of 0 – 600 µmol/mL in 1 mL of milliQ water. The solution was then heated at 60^°^C for 10 min and stored at room temperature (RT). 5FU (Sigma-Aldrich, Australia) was added to the copper-polyethylenimine complex and incubated at 60ºC for 1 h to form the PEI-Cu-5FU complex. The final concentration of 5FU was 50 mM.

Stealth neutral liposomes were prepared by dissolving DSPC:cholesterol:PEG2000-DSPE (ratio 65:30:5) (Avanti Polar Lipids, Alabaster, Alabama, USA) in chloroform (Sigma Aldrich, Australia). The chloroform was removed under vacuum (18 h) to form lipid films. The film was hydrated by adding 5FU solution to create a neutral liposomal (NL) 5FU dispersion, or by adding the solution of PEI-Cu-5FU to create a NL-PEI-Cu-5FU dispersion.

Cationic liposomes were prepared by dissolving DSPC:DC-6-14:cholesterol:PEG2000-DSPE (ratio 30:35:30:5) (Sogo Pharmaceuticals, Tokyo, Japan) in chloroform. The chloroform was removed under vacuum (18 h) to form lipid films. The film was hydrated by the PEI-Cu-5FU solution to create a cationic liposomal (CL) PEI-Cu-5FU dispersion.

All liposomal dispersions were then extruded through a 100 nm polycarbonate Lipex membrane (Northern Lipids, Burnaby, Canada), and centrifuged to remove the supernatant. The remaining liposomal pellets were re-dispersed prior to intravenous administration.

### Experimental design

All experiments were performed with inbred male Sprague Dawley (SD) rats (SA Pathology, Adelaide, Australia), weighing between 150 g and 170 g. Approval for conduct of the experiments was provided by SA Pathology Animal Ethics Committee. Three rats were in each housing in an animal facility regulated at 22 ± 1ºC and subject to a 14:10 h light-dark cycle.

Rats were randomly assigned to groups; saline (n=6), 5FU (n=6), 5FU-PEI-Cu (n=6), NL-5FU (n=5), NL-PEI-Cu-5FU (n=5), and CL-PEI-Cu-5FU (n=5) groups. All treatments were administered intravenously via the tail vein once daily for five days, with each formulation containing 5FU administered in phosphate buffer (pH 7.4, 1 mM) at a dose of 10 mg/kg. Rats were killed by exsanguination and cervical dislocation two days after the final injection while under anaesthesia with 3% isofluorane in 100% O_2_.

The small intestine (pyloric sphincter to ileocaecal sphincter) was dissected out and flushed with chilled, sterile saline. Samples for histological analyses (1 cm) were taken at approximately 50% of the length (jejunum).

### Histopathology

The histological procedure was described previously [12]. Briefly, routine haematoxylin and eosin staining was used to observe histopathological changes in jejunum and colon samples. Scoring was analysed based on 10 fields of view. Individual parameters for villous fusion, villous blunting, crypt ablation, and inflammatory infiltrate were graded compared to normal tissue from 0 to 3. 0 = no change from standard histological structure, 1 = up to one third of tissue affected, 2 = between one and two thirds of tissue affected, and 3 = greater than two thirds of tissue affected [12].

### Analysis of goblet cells

The procedure and scoring was described previously [12]. Briefly, Alcian Blue/PAS staining was used to analyse goblet cells and mucins. Sections were dewaxed and rehydrated in xylene and graded series of alcohol respectively. Sections were then stained in Alcian Blue (1% Alcian Blue 8GX (CI 74240, Sigma-Aldrich) in 3% glacial acetic acid (Sigma-Aldrich)), rinsed in distilled water, and oxidised with 1% periodic acid prior to washing. Sections were then treated in Schiff’s reagent for 15 min, washed, dehydrated, cleared and mounted.

Total goblet cells, including cavitated goblet cells (apical indentation into the intracellular store of mucus granules, indicative of accelerated mucus secretion by compound exocytosis) [13] and paired goblet cells (adjacent goblet cells), were counted for a minimum of fifteen intact villi and/or crypts per section. Goblet cell composition was also analysed, with blue staining, magenta staining, and purple staining demonstrating acidic, neutral, and mixed mucins respectively [12].

### Immunohistochemistry

Experimental design was described previously [12]. Briefly, the enteric nervous system (ENS) was investigated using immunohistochemistry. The anti-S-100 antibody (Dako, Denmark) has previously been shown to positively stain neurons and glial cells of the peripheral nervous system (PNS) (manufacturer data sheet). Stained cells were counted over a 1 mm length of tissue, with large enteric ganglia (groups of four or more stained cells), and a total of all stained cells were recorded. The location of the positively stained cells was also noted. Staining intensity was also graded as a semiquantitative measure of the level of expression, where 0= negative; 1= weak; 2= moderate; 3= strong; 4=very intense, based on a previously validated technique [14].

### Statistical analyses

Means between each group were compared using the Kruskal-Wallis test with a secondary Dunn’s comparative test for multiple comparisons. (GraphPad Prism 5.0, Graphpad software, Califorina, USA). Asterix denotes significance at p <0.05. Effect size for clinical significance was analysed with Cohen’s D tests comparing rats receiving nanoparticles with rats receiving 5FU only. The likely clinical significance of the effect was considered small if d > 0.20, moderate if d > 0.50, and large if d > 0.80 [15].

## Results

### Liposomal formulation maintains histological structure and increases inflammatory infiltrate in the small intestine

Histological changes were observed along the GIT following 5FU administration or PEI-Cu administration. In the jejunum, villous fusion significantly increased in PEI-Cu-5FU (formulation vehicle) and NL-PEI-Cu-5FU rats compared with saline control rats (p = 0.002). Inflammatory infiltrate also significantly increased in PEI-Cu-5FU and NL-PEI-Cu-5FU rats compared with the saline control rats (p = 0.0011). Furthermore, inflammatory infiltrate was also significantly higher in the PEI-Cu-5FU rats compared with the 5FU rats (p = 0.0011, d = -5.5, large effect). Villous blunting and crypt ablation did not change significantly following 5FU treatments (Figure 1).

**Figure 1.**
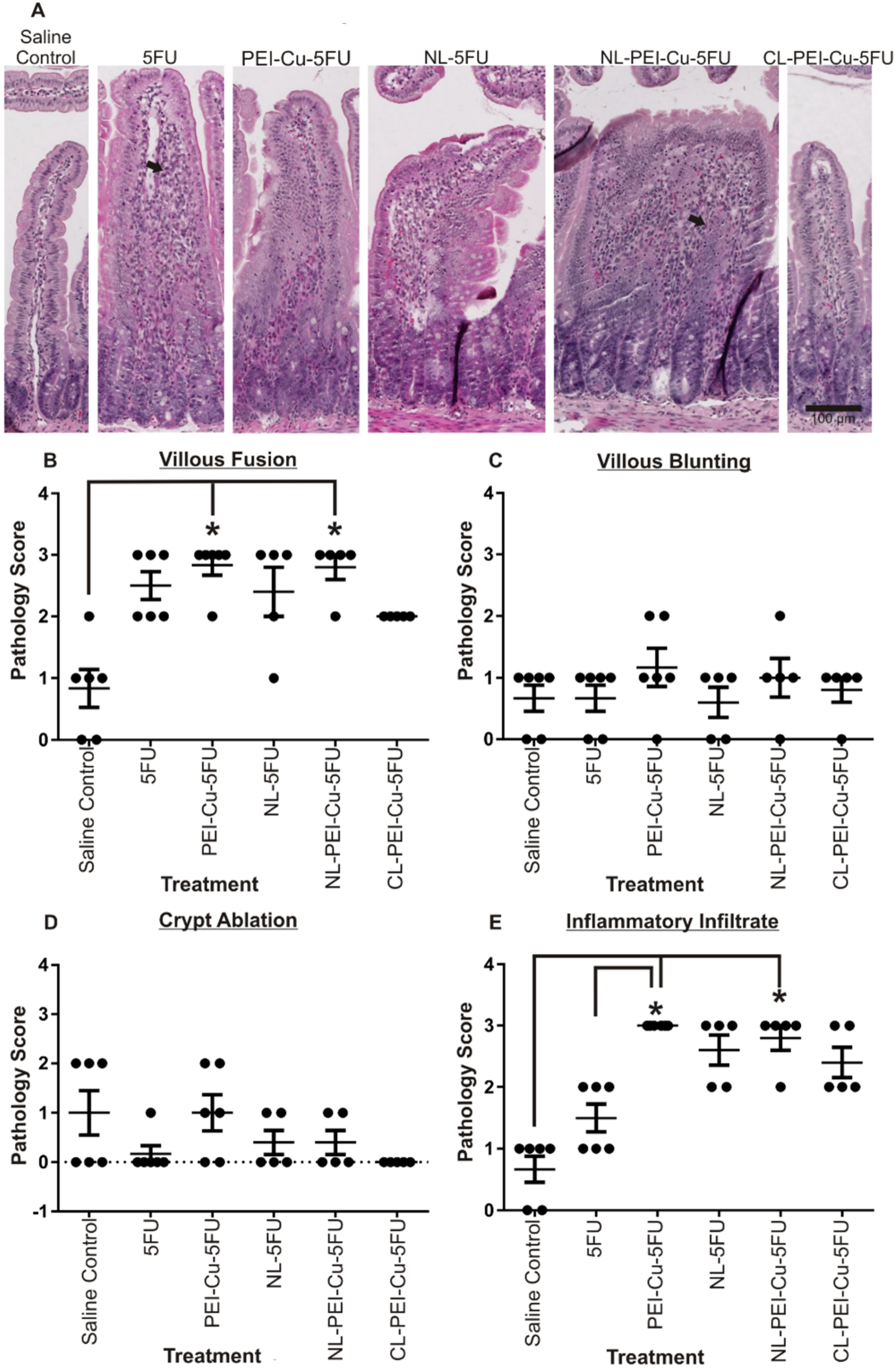
*Histopathology*. A. In the jejunum villous blunting increased in PEI-Cu-5FU rats and NL-PEI-Cu-5FU rats compared to the saline control (p=0.002), and inflammatory infiltrate increased in PEI-Cu-5FU rats compared to the 5FU rats (p=0.0011). In the 5FU section the arrow highlights inflammatory infiltrate; in the NL-PEI-Cu-5FU section the arrow highlights villous fusion. (Original Magnification 20x) B. Villous fusion scores in jejunum. C. Villous blunting scores in the jejunum D. Crypt ablation scores in the jejunum E. Inflammatory Infiltrate scores in the jejunum.

### Liposomal formulation affects goblet cells in the in the small intestine

#### Liposomal formulation affects goblet cell number

In the jejunum of saline control rats, the total number of goblet cells was 20.5 ± 1.8 (mean ± SEM) per villus and 7.8 ± 0.8 per crypt. These numbers were lowest in 5FU treated rats (9.3 ± 1.4 per villus and 4.2 ± 0.8 per crypt). The total number of goblet cells in PEI-Cu-5FU (formulation vehicle) treated rats was 12.9 ± 1.4 per villus (d = -1.03, large effect) and 5.7 ± 1.0 per crypt (d = -0.73, medium effect). In the groups treated with the liposomal formulations, the total number of goblet cells per villus was 16.5 ± 3.7 in the NL-PEI-Cu-5FU treated rats (d = -1.32, large effect), and 19.28 ± 2.9 in NL-5FU treated rats (d = -2.15, large effect). In the crypts the total number of goblet cells was lowest in the CL-PEI-Cu-5FU rats (5.8 ± 0.9, d = -0.82, large effect) and highest in the NL-PEI-Cu-5FU rats (8.4 ± 2.1, d = -1.38, large effect). The mean ranks of the total number of goblet cells per villus deviated significantly across all groups (p = 0.0115) (Figure 2).

**Figure 2.**
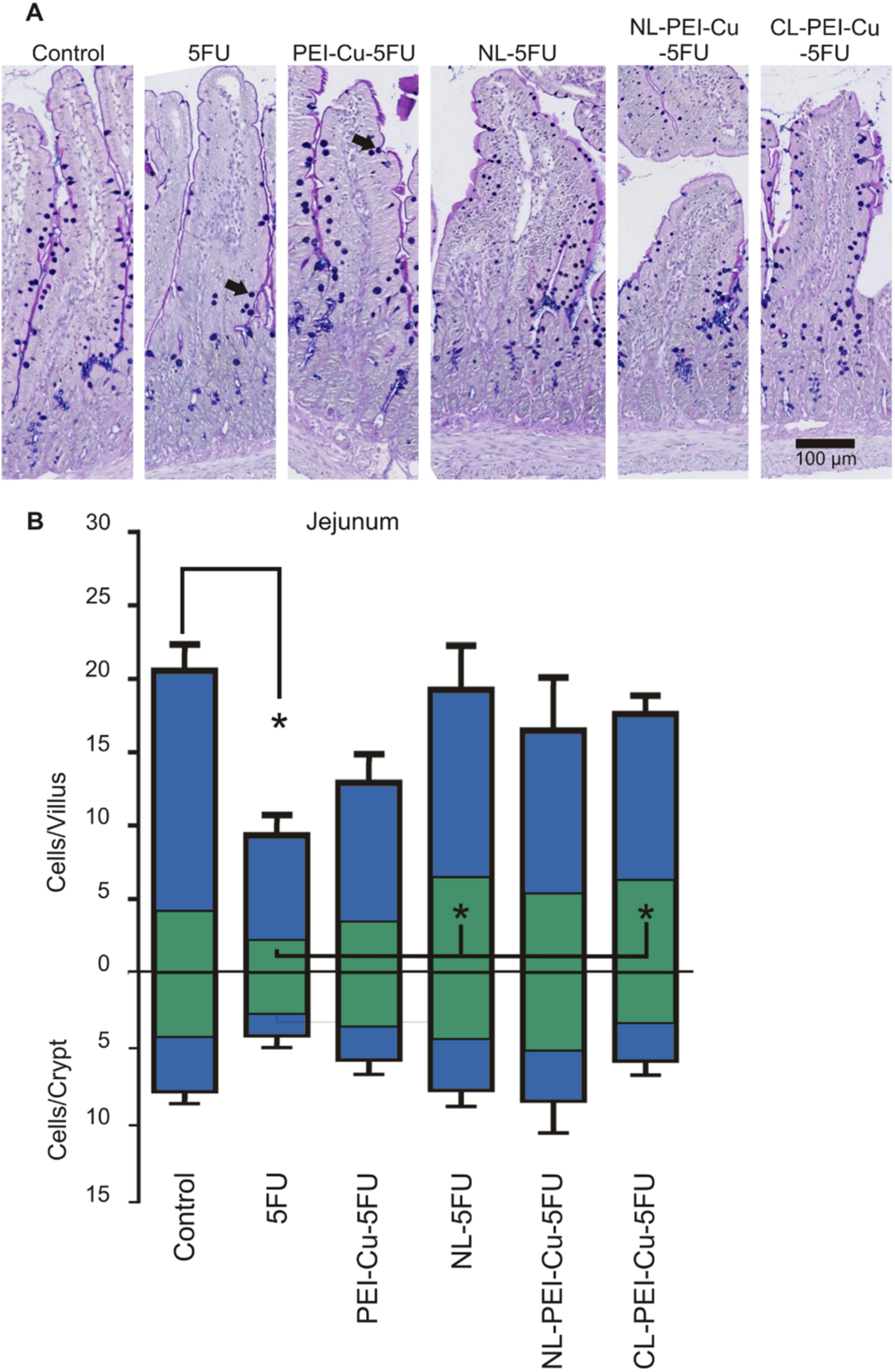
*Alcian Blue-PAS staining* A. Villous and crypts of jejunum. (Original Magnification 20x). B. Goblet cell counts of intact and cavitated goblet cells in the jejunum. Significant decrease in 5FU villous total goblet cell numbers compared to control (p<0.05). Significant increase in NL-5FU and CL PEI-Cu-5FU villous cavitated goblet cell numbers compared to 5FU (p<0.05).

#### Liposomal formulation affects the percentage of cavitated goblet cells

Cavitated goblet cells (expressed as a percentage of total goblet cells) in the jejunums of saline control and 5FU rats was 20.2 ± 4.2 and 24.5 ± 3.6, respectively, and in the crypts 52.1 ± 8.4 and 73.8 ± 5.0, respectively. The percentages of cavitated goblet cells for the formulation vehicle PEI-Cu-5FU group was 25.2 ± 2.2 in the villi (d = -0.10), and 66.4 ± 4.6 in the crypts (d = 0.63, medium effect). The percentages of cavitated goblet cells in the villi were similar in the NL-PEI-Cu-5FU, NL-5FU, and CL-PEI-Cu-5FU rats. These were 38.7 ± 10.6 (d = -0.95, large effect), 35.8 ± 8.0 (d =-0.91, large effect), and 35.2 ± 3.5, (d = -1.31, large effect), respectively. In the crypts, the percentages of cavitated goblet cells were also similar in the NL-PEI-Cu-5FU, NL-5FU, and CL-PEI-Cu-5FU rats. These were 58.8 ± 4.0 (d = 1.00, large effect), 57.1 ± 10.6 (d = 1.40, large effect), and 59.4 ± 4.6 (d = 1.27, large effect), respectively. The mean ranks of the percentages of cavitated cells did not deviate significantly (Figure 2).

#### Liposomal formulation affects the percentage of paired goblet cells

Paired goblet cells (expressed as a percentage of total goblet cells) in the villi of jejunums of saline control and 5FU rats were 27.5 ± 9.2 and 17.3 ± 7.4, respectively. The percentage of paired goblet cells for the formulation vehicle PEI-Cu-5FU rats was 46.5 ± 10.0 (d = -1.37, large effect). The percentages of paired goblet cells in the villi of the NL-PEI-Cu-5FU, NL-5FU, and CL-PEI-Cu-5FU groups were 24.1 ± 10.0 (d = -1.07, large effect), 38.3 ± 10.5 (d = -0.36, small effect), and 30.7 ± 7.7 (d = -0.75, medium effect), respectively. The mean ranks of the percentages of paired cells did not deviate significantly (Supplementary Figure 1).

### Immuno-labelling of S100 positive cells

S-100 positive cells (neurons and glial cells) were located in the submucosal and myenteric plexuses of the jejunum. S100 positive cells were also observed in the circular layer of the muscularis externa. The staining intensity of S100 positive cells and enteric ganglia in the submucosal and myenteric plexuses of the jejunum increased following 5FU, PEI-Cu-5FU, NL-PEI-Cu-5FU, NL-5FU, and CL-PEI-Cu-5FU compared to the saline control, with a significant difference in mean ranks of the intensity of staining in the myenteric plexus (p=0.0471) (Figure 3).

**Figure 3.**
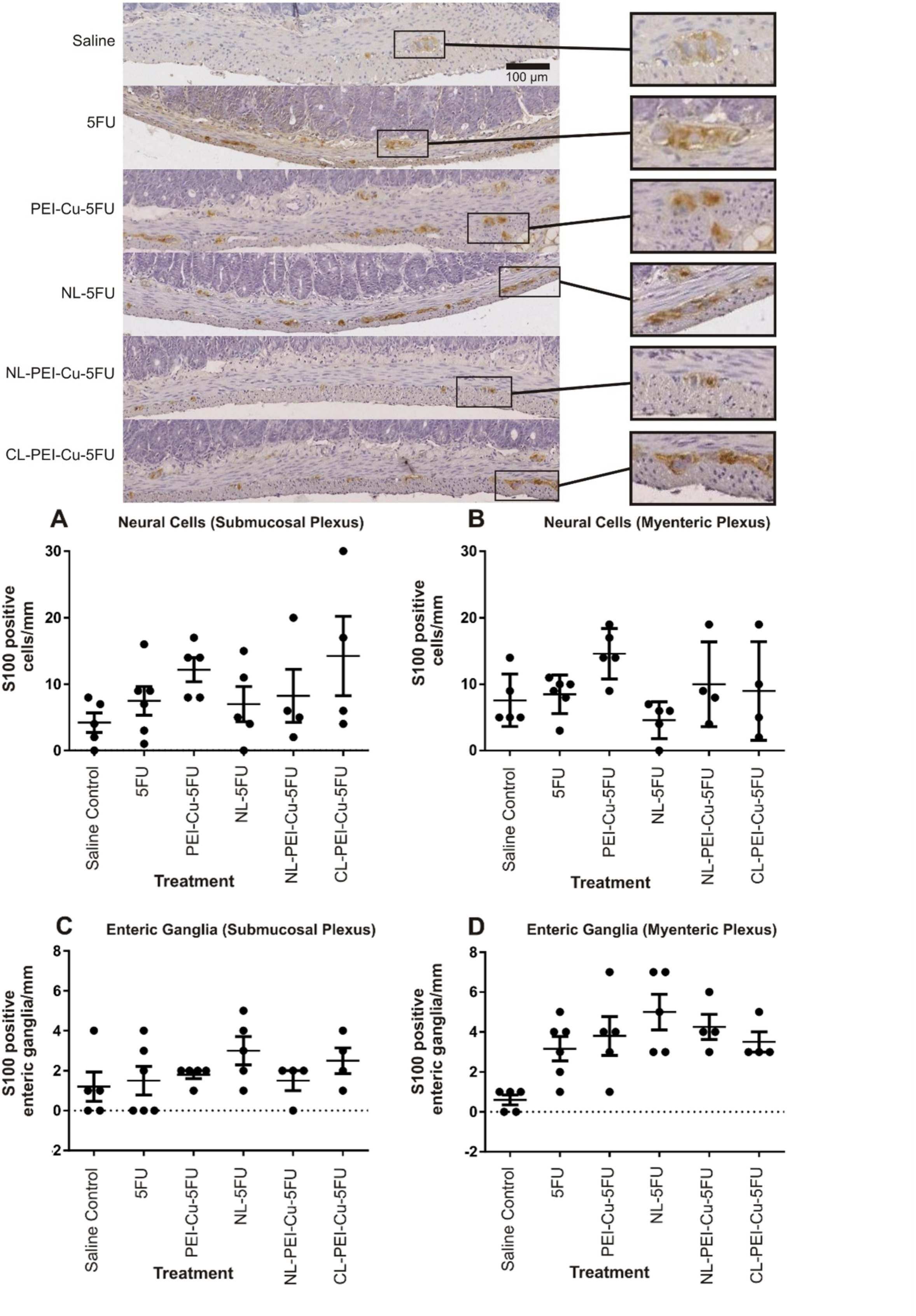
*(Top) Immunohistochemistry of S100*. Positive staining of cells in the jejunum, inset boxes highlight enteric ganglia in the respective sections. (Original Magnification 20X). (*Bottom) S100 cell counts* A. Positive staining of neural cells in jejunum submucosal plexus. B. Positive neural cells in the jejunum myenteric plexus. C. Positive enteric ganglia in the jejunum submucosal plexus. D. Positive enteric ganglia in the jejunum myenteric plexus.

The staining intensities of S-100 positive cells in the jejunum was 2.0 ± 0.4 and 3.1 ± 0.4 in the submucosal plexus, and 2.3 ± 0.4 and 3.7 ± 0.2 in the myenteric plexus, for saline and 5FU rats, respectively. No changes in intensity were observed between 5FU and other 5FU formulations (Supplementary Figure 2).

### Quantification of S100 positive cells

#### S100 positive cells

The numbers of S100 positive cells in the submucosal plexus in the jejunum of saline and 5FU rats were 4.2 ± 1.5 and 7.5 ± 2.2 cells per millimetre (mm), respectively. The mean ranks of the number of neural cells per mm in the submucosal plexus did not deviate significantly. The numbers of neural cells in the myenteric plexus in the jejunum of saline and 5FU rats were 7.6 ± 1.8 and 8.5 ± 1.2 cells per mm, respectively. This number was highest in the PEI-Cu-5FU rats (14.6 ± 1.7, d = -1.83, large effect). However, the mean ranks of neural cells per mm in the myenteric plexus did not deviate significantly (Figure 3).

#### Enteric ganglia

The numbers of enteric ganglia in the submucosal plexus in the jejunum of saline and 5FU rats was 1.2 ± 0.7 and 1.5 ± 0.7 enteric ganglia per mm, respectively. Numbers of enteric ganglia were highest in the NL-5FU rats (3.0 ± 0.7 per mm, d = -1.06, large effect). However, the mean ranks of the number of enteric ganglia per mm in the submucosal plexus did not deviate significantly (Figure 3).

The numbers of enteric ganglia in the myenteric plexus of the jejunum in saline and 5FU rats was 0.6 ± 0.2 and 3.2 ± 0.6, enteric ganglia per mm, respectively. There was a significant increase in enteric ganglia in the NL-5FU rats compared with saline rats to 5.0 ± 0.9 (p<0.05). The mean ranks of the number of enteric ganglia per mm in the myenteric plexus deviated significantly across groups (p = 0.0212) (Figure 3).

### Non-parametric regression

Non-parametric regression analysis demonstrated a strong correlation in the jejunum between villous fusion, and inflammatory infiltrate, intact goblet cells per villus, the percentage of paired goblet cells per villus, myenteric plexus staining intensity and submucosal staining intensity. Furthermore, strong correlations occurred between inflammatory infiltrate, and the number of enteric ganglia in the myenteric plexus and staining intensity in the myenteric plexus (Supplementary Table 1).

## Discussion

5FU, a cytotoxic chemotherapy agent, has been shown to cause mucositis with the most severe damage occurring in the small intestine [16]. Liposomes and PEI-Cu offer an increased plasma circulation time of 5FU, causing an increase in cytotoxicity and antineoplastic effect [1]. The effects of the increased circulation time on mucositis and other toxic side effects are largely unknown. This study has demonstrated specific differences in the 5FU-induced damage in the jejunum with the use of 5FU carrier molecules PEI-Cu-5FU.

Previous studies have focused on increasing the cytotoxic effects on cancerous cells while broadly looking at toxicity [1-4, 17]. Specifically, 5FU attached to PEI-Cu in liposomes had a greater anti-tumour effect than 5FU alone and resulted in a similar percentage of weight change [1]. Cosco et al. (2009) provided supporting results with an increase in the cytotoxicity of 5FU when loaded with polyetheylene-glycol coated bola-niosomes in MCF-7 and T47D breast cancer cell lines, and *in vivo* in a xenograft of MCF-7 tumours [17]. No study has yet determined the effect of liposomes on the gut in relation to the toxicities associated with 5FU use. However, increased 5FU cytotoxicity may result in increased mucositis.

This study has shown histological evidence of mucositis in the 5FU group of rats, with inflammatory infiltrate and villous fusion significantly (P ≤ 0.05) increased in the mucosa of 5FU treated rats compared to the saline group, consistent with previous studies [10, 18, 19]. Mucin composition in the 5FU group compared to the saline control group was also similar compared to previous studies of 5FU-induced mucositis, with no significant change in total goblet cell numbers or percentage of cavitated goblet cells in the jejunum [10].

This study investigated differences in the histopathology based on the different carrier molecule formulations, with inflammatory infiltrate significantly increasing (p<0.05) in PEI-Cu-5FU treated rats compared to 5FU. This increase may be caused by an increase in circulation time in PEI-Cu-5FU treated rats compared to 5FU treated rats [1]. Furthermore, increased circulation time would also explain the slight but not significant rise of inflammatory cells in NL-5FU, NL-PEI-Cu-5FU and CL-PEI-Cu-5FU [1].

Mucin stores in goblet cells were also shown to differ between carrier molecule formulations. An increase in the percentage of cavitated goblet cells in NL-5FU and CL-PEI-Cu-5FU rats compared to 5FU-treated rats was shown, suggesting mucin secretion may be increasing with these formulations. Furthermore, the percentage of paired goblet cells trends towards an increase in the villi of PEI-Cu 5FU rats compared to the 5FU rats with a (d = -1.37) large effect size (not significant). Goblet cell precursor cells cannot create two adjacent goblet cells, resulting from notch signalling [20]. Therefore, adjacent goblet cells are indicative that enterocytes between have died and suggests increased enterocyte cell death in the PEI-Cu-5FU rats compared to 5FU rats.

The number of neural cells in the myenteric plexus of NL-5FU rats trended towards a decrease compared to 5FU rats with a (d = -1.06) large effect size (not significant). Of these, the positive cells that are neurons could be sensory neurons, interneurons, or motor neurons. Sensory neuron loss would affect the ability to detect chemical and mechanical stimuli, interneurons would affect transmission of data to motor neurons, and motor neuron loss may affect smooth muscle contraction and distension, and secretion. The reduced number of neural cells following NL-5FU, and whether they are sensory, inter-, or motor neurons is likely to result in reduced neural stimulation of smooth muscle and secretory cells. However, the findings in this study suggest that the neural loss is likely not associated with goblet cell secretion or mucin stores. Conversely, an increase in S100 positive neurons in 5FU-PEI-Cu rats compared to saline and 5FU rats was observed as well as a correlation between increased inflammation and S100 positive cell numbers. This is unlikely to occur in adults and would suggest that the increase in S100-positive neurons and glial cells is not due to an increase of cells, but more likely due to an increased production of S100 protein.

Based on these findings, it is worthwhile investigating these and other carrier molecule formulations of 5FU and other cytotoxic agents further. These formulations showed varied but small changes in cytotoxicity in the gut suggesting increased tumour cytotoxicity without or with limited gastrointestinal toxicity. Due to this duality these formulations are likely to become more widely used in the future. However, investigations into the specific effects of these formulations on enteric neural cells is also warranted, in particular functional and electrophysiological studies with enteric neurons to see the differences between these formulations and conventional cytotoxic agents, and the impact they may have on chemotherapy-induced mucositis and other related toxicities.

In conclusion, this study showed, for the first time, differences in gastrointestinal toxicity and damage between different formulations of molecular carriers for 5FU, with preliminary suggestions that intestinal parameters may not be negatively impacted by these formulations that increase cytotoxicity. This preliminary data indicates the need for more in-depth studies investigating the specific effects of carrier molecules for cytotoxic agents due to the severe dose limiting nature of gastrointestinal toxicities compared to conventional chemotherapy regimens.

### Statements and Declarations

On behalf of all authors, the corresponding author states that there is no conflict of interest.

The datasets generated during and/or analysed during the current study are available from the corresponding author on reasonable request

## Acknowledgements

This study was funded by National Health and Medical Research Council funding (1016696)

## Notes

**<u>Statements and Declarations</u>** Conflict of Interest: The authors declare that they have no conflict of interest.

### Competing Interest Statement

The authors have declared no competing interest.

## References

1. Thomas, A.M., et al., Development of a liposomal nanoparticle formulation of 5-fluorouracil for parenteral administration: formulation design, pharmacokinetics and efficacy. J Control Release, 2011. 150(2): p. 212–9.

2. Ahmad, N., et al., A comparative ex vivo permeation evaluation of a novel 5-Fluorocuracil nanoemulsion-gel by topically applied in the different excised rat, goat, and cow skin. Saudi Journal of Biological Sciences, 2020. 27(4): p. 1024–1040.

3. Patras, L., et al., Liposomal prednisolone phosphate potentiates the antitumor activity of liposomal 5-fluorouracil in C26 murine colon carcinoma in vivo. Cancer Biol Ther, 2017. 18(8): p. 616–626.

4. Dupertuis, Y.M., et al., Antitumor Effect of 5-Fluorouracil-Loaded Liposomes Containing n-3 Polyunsaturated Fatty Acids in Two Different Colorectal Cancer Cell Lines. Aaps Pharmscitech, 2021. 22(1): p. 10.

5. Yaman, U., et al., Surface modified nanoliposome formulations provide sustained release for 5-FU and increase cytotoxicity on A431 cell line. Pharmaceutical Development and Technology, 2020. 25(10): p. 1192–1203.

6. Scavo, M.P., et al., Effectiveness of a Controlled 5-FU Delivery Based on FZD10 Antibody-Conjugated Liposomes in Colorectal Cancer In vitro Models. Pharmaceutics, 2020. 12(7): p. 19.

7. He, W., et al., Dimeric Artesunate–Phosphatidylcholine-Based Liposomes for Irinotecan Delivery as a Combination Therapy Approach. Molecular Pharmaceutics, 2021. 18(10): p. 3862–3870.

8. Abu Lila, A.S., et al., Oxaliplatin encapsulated in PEG-coated cationic liposomes induces significant tumor growth suppression via a dual-targeting approach in a murine solid tumor model. Journal of Controlled Release, 2009. 137(1): p. 8–14.

9. Biswas, S., et al., Liposomes loaded with paclitaxel and modified with novel triphenylphosphonium-PEG-PE conjugate possess low toxicity, target mitochondria and demonstrate enhanced antitumor effects in vitro and in vivo. J Control Release, 2012. 159(3): p. 393–402.

10. Stringer, A.M., et al., Gastrointestinal microflora and mucins may play a critical role in the development of 5-Fluorouracil-induced gastrointestinal mucositis. Exp Biol Med (Maywood), 2009. 234(4): p. 430–41.

11. Costa, D.V.S., et al., 5-Fluorouracil Induces Enteric Neuron Death and Glial Activation During Intestinal Mucositis via a S100B-RAGE-NFκB-Dependent Pathway. Scientific Reports, 2019. 9(1): p. 665.

12. Thorpe, D., et al., Irinotecan-Induced Mucositis Is Associated with Goblet Cell Dysregulation and Neural Cell Damage in a Tumour Bearing DA Rat Model. Pathology and Oncology Research, 2020. 26(2): p. 955–965.

13. Barcelo, A., et al., Mucin secretion is modulated by luminal factors in the isolated vascularly perfused rat colon. Gut, 2000. 46(2): p. 218–24.

14. Bowen, J.M., et al., Cytotoxic chemotherapy upregulates pro-apoptotic Bax and Bak in the small intestine of rats and humans. Pathology, 2005. 37(1): p. 56–62.

15. Cohen, J., Statistical Power Analysis for the Behavioral Sciences. 2013: Taylor & Francis.

16. Fata, F., et al., 5-fluorouracil-induced small bowel toxicity in patients with colorectal carcinoma. Cancer, 1999. 86(7): p. 1129–34.

17. Cosco, D., et al., Novel PEG-coated niosomes based on bola-surfactant as drug carriers for 5-fluorouracil. Biomed Microdevices, 2009. 11(5): p. 1115–25.

18. Saegusa, Y., et al., Changes in the mucus barrier of the rat during 5-fluorouracil-induced gastrointestinal mucositis. Scand J Gastroenterol, 2008. 43(1): p. 59–65.

19. Logan, R.M., et al., Is the pathobiology of chemotherapy-induced alimentary tract mucositis influenced by the type of mucotoxic drug administered? Cancer Chemother Pharmacol, 2009. 63(2): p. 239–51.

20. Sancho, R., et al., Fbw7 repression by hes5 creates a feedback loop that modulates Notch-mediated intestinal and neural stem cell fate decisions. PLoS Biol, 2013. 11(6): p. e1001586.

