## Supplementary figures and images for "Different liposomal formulations of 5-Fluorouracil result in variations to gastrointestinal toxicities"

### Supplemental figure 1

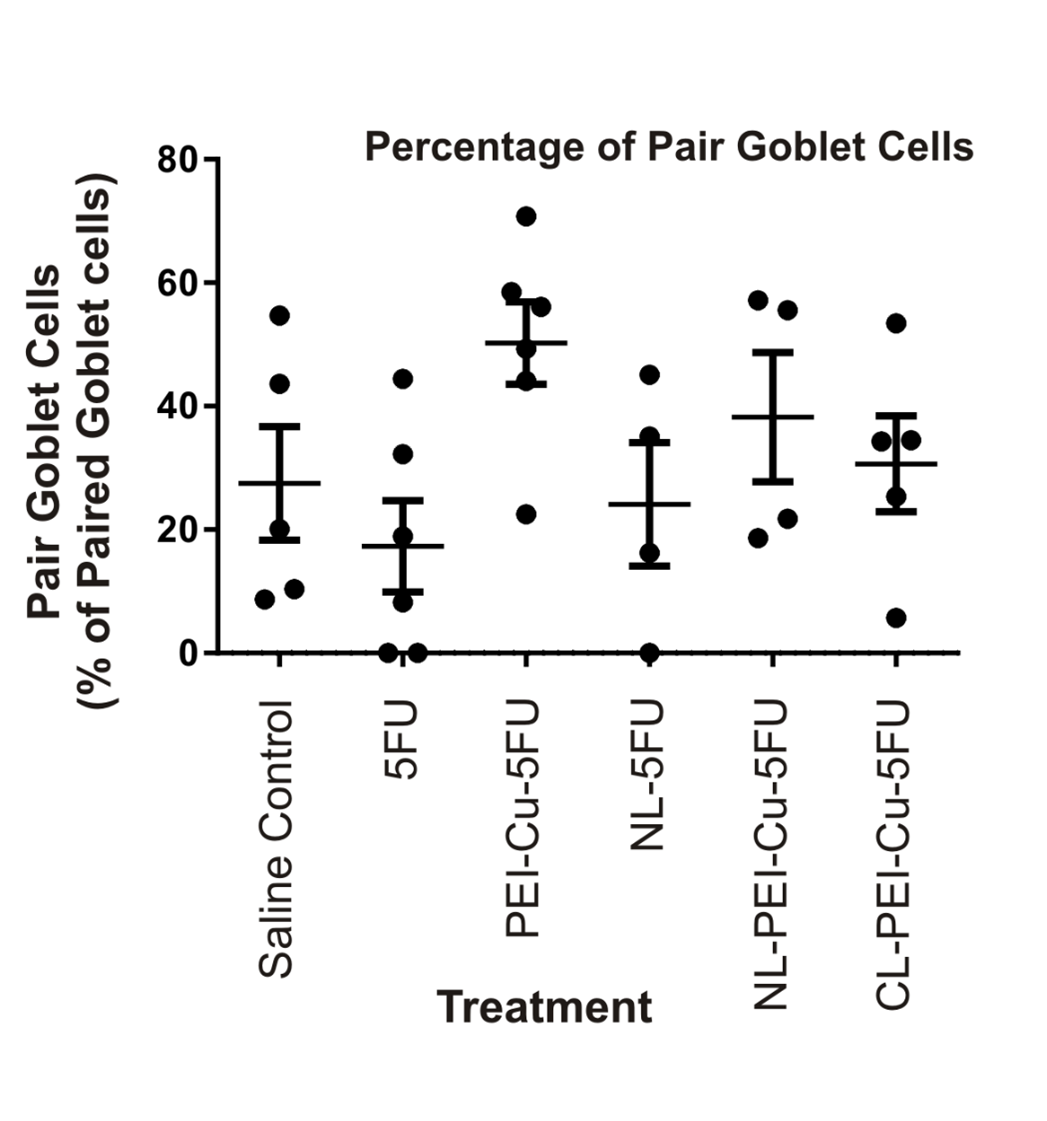

### Supplemental figure 2

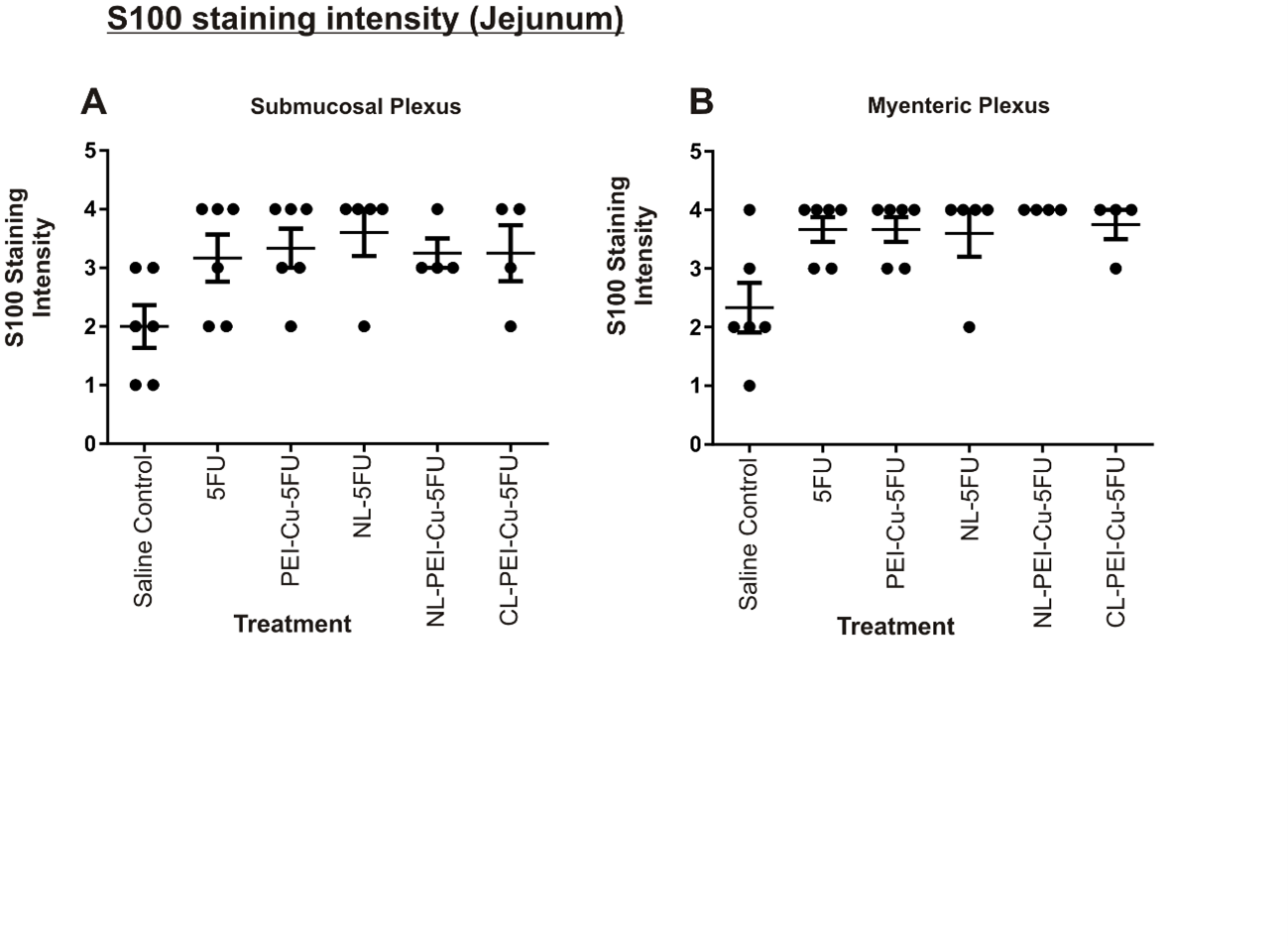
