## Supplemental Table 1 for "Different liposomal formulations of 5-Fluorouracil result in variations to gastrointestinal toxicities"

**Supplementary Table 1** Non parametric regression.

| **Gut Region** | **Dependent Variable** | **Independent Variable** | **Coefficient** | **95% CI** | **Pseudo R²** |
| --- | --- | --- | --- | --- | --- |
| **Jejunum** | Villous Fusion | Inflammatory Infiltrate | 1.00 | 0.86  1.14 | 0.52 |
| **Jejunum** | Villous Fusion | Intact Goblet Cells (Villus) | -0.09 | -0.17  -0.01 | 0.16 |
| **Jejunum** | Villous Fusion | % Paired Goblet Cells (Villus) | 0.02 | 0.00  0.03 | 0.01 |
| **Jejunum** | Villous Fusion | S100 Staining Intensity (Myenteric Plexus) | 1.00 | 0.61  1.39 | 0.27 |
| **Jejunum** | Villous Fusion | S100 Staining Intensity (Submucosal Plexus) | 1.00 | 0.54  1.46 | 0.18 |
| **Jejunum** | Inflammatory Infiltrate | Enteric Ganglia (Myenteric Plexus) | 0.33 | 0.09  0.58 | 0.25 |
| **Jejunum** | Inflammatory Infiltrate | S100 Staining Intensity (Myenteric Plexus) | 1.00 | 0.61  1.39 | 0.24 |
| **Jejunum** | Intact Goblet Cells (Villus) | Cavitated Goblet Cells (Villus) | 1.36 | 0.09  2.62 | 0.11 |
| **Jejunum** | Intact Goblet Cells (Villus) | % Cavitated Goblet Cells (Villus) | -0.26 | -0.43  -0.09 | 0.20 |
| **Jejunum** | Intact Goblet Cells (Villus) | Intact Goblet Cells (Crypt) | 1.77 | 0.11  3.43 | 0.10 |
| **Jejunum** | Cavitated Goblet Cells (Villus) | % Cavitated Goblet Cells (Villus) | 0.09 | 0.02  0.17 | 0.21 |
| **Jejunum** | % Cavitated Goblet Cells (Villus) | % Paired Goblet Cells (Villus) | 0.18 | 0.01  0.35 | 0.12 |
| **Jejunum** | Intact Goblet Cells (Crypt) | Cavitated Goblet Cells (Crypt) | 0.37 | 0.04 0.71 | 0.10 |
| **Jejunum** | Intact Goblet Cells (Crypt) | % Cavitated Goblet Cells (Crypt) | -0.07 | -0.09  -0.04 | 0.40 |
| **Jejunum** | S100 Nuclei (Myenteric Plexus) | S100 Enteric Ganglia (Submucosal Plexus) | 4.00 | 0.02  7.98 | 0.09 |
| **Jejunum** | S100 Nuclei (Submucosal Plexus) | S100 Enteric Ganglia (Submucosal Plexus) | 3.00 | -0.00  6.00 | 0.13 |
| **Jejunum** | S100 Neural Cells (Submucosal Plexus) | S100 Neural Cells (Myenteric Plexus) | 0.78 | 0.31  1.24 | 0.27 |
| **Jejunum** | S100 Enteric Ganglia (Submucosal Plexus) | S100 Enteric Ganglia (Myenteric Plexus) | 0.33 | 0.05  0.62 | 0.18 |
| **Jejunum** | S100 Enteric Ganglia (Submucosal Plexus) | S100 Staining Intensity (Submucosal Plexus) | 1.00 | 0.12  1.88 | 0.10 |
| **Jejunum** | S100 Enteric Ganglia (Myenteric Plexus) | S100 Staining Intensity (Myenteric Plexus) | 2.00 | 0.86  3.14 | 0.21 |
